# Genome-wide mediation analysis: bridging the divide between genotype and phenotype via transcriptomic data in maize

**DOI:** 10.1101/2021.04.09.439238

**Authors:** Zhikai Yang, Gen Xu, Qi Zhang, Toshihiro Obata, Jinliang Yang

**Affiliations:** Department of Agronomy and Horticulture, University of Nebraska-Lincoln, Lincoln, NE 68588, USA; Center for Plant Science Innovation, University of Nebraska-Lincoln, Lincoln, NE 68583, USA; Department of Mathematics and Statistics, University of New Hampshire, Durham, NH 03824, USA; Department of Biochemistry, University of Nebraska-Lincoln, Lincoln, NE 68583, USA

## Abstract

Mapping genotype to phenotype is an essential topic in genetics and genomics research. As the Omics data become increasingly available, genome-wide association study (GWAS) has been widely applied to establish the relationship between genotype and phenotype. However, signals detected by GWAS usually span broad genomic regions with many underneath candidate genes, making it challenging to interpret and validate the molecular functions of the candidate genes. Under the context of genetics research, we hypothesized a causal chain from genotype to phenotype partially mediated by intermediate molecular processes, i.e., gene expression. To test this hypothesis, we applied the high dimensional mediation analysis, a class of causal inference method with an assumed causal chain from the exposure to the mediator to the outcome, and implemented it to the maize diversity panel (N=280 lines). Using 40 publicly available agronomic traits, 66 newly generated metabolic traits, and published RNA-seq data from seven different tissues, we detected N=736 unique mediating genes, explaining an average of 12.7% phenotypic variance due to mediation. Noticeably, 83/736 (11%) genes were identified in mediating more than one trait, suggesting the prevalence of pleiotropic mediating effects. Among those pleiotropic mediators, benzoxazinone synthesis 13 (*Bx13*), a well-characterized gene encoding a 2-oxoglutarate-dependent dioxygenase, was identified mediating 40 agronomic and metabolic traits in different tissues. Further genetic and genomic analyses of the *Bx13* and adjacent mediating genes suggested a 3D co-regulation modulation likely affect their expression levels and eventually lead to phenotypic consequences. Our results suggested the genome-wide mediation analysis is a powerful tool to integrate Omics data in providing causal inference to connect genotype to phenotype.

## INTRODUCTION

By exploiting the genetic potential of a few genes, Green Revolution and following technical innovations have pushed the crop yield to their limits [1, 2]. As the world population size marches toward 9 billion, the crop yield is expected to improve by 50% to meet the increasing calorie demand by 2050 [3]. To maximize crop yield, it is crucial to further exploit the genetic potential and establish the relationship between agronomically important phenotypes with genotypes (G2P). These crop yield and other agronomically essential phenotypes, however, are quantitative traits usually determined by many genes. The underlying genes are likely being modulated by interconnected regulatory networks and involved in complex molecular pathways. As Omics datasets become increasingly available, integrating these Omics datasets to connect G2P provides a promising opportunity.

Recently emerged technologies leverages the linkage disequilibrium (LD) between true causal variants and the single nucleotde polymorphism (SNP) markers to identify trait-marker association (i.e., genome-wide association study, or GWAS) or predict the individual’s phenotypic performance (i.e., genomic selection, or GS) [4, 5]. However, the essence of association (but not causation) for both of these methods prevents their in-depth applications as the associated markers may be uncoupled with the causal variants in a different population. From a basic research point of view, GWAS usually detects SNPs within long blocks of genomic regions, in which many candidate genes might reside, challenging the downstream gene functional studies. Even if the causal SNPs can be identified, the genes underlying the phenotypes are not necessarily the ones closest to the SNPs. Instead, the GWAS signals may be linked with causal genes through long-range chromatin interactions or distal gene regulation [6]. To overcome such limitations, practical methods to bridge G2P by elucidating the intermediate molecular mechanisms require to be developed. In an attempt to dissect the molecular mechanisms in controlling the phenotypic variation, a number of studies have been performed by considering transcriptomics data, such as the eQTL analysis, transcriptome-wide association study, and coexpression network-based gene prioritization [7–9]. These studies provided insights into the possible causes of gene expression variation and their functional consequences. Nevertheless, they did not model genotype, gene expression, and phenotype altogether.

Mediation analysis, as one of the causal inference methods, has the potential to build a causal connection between marker and phenotype by including a third hypothetical variable, e.g., gene expression. Mediation analysis is carried out based on an assumed causal chain that was first proposed by Baron and Kenny [10]. Recently, a high dimensional mediation model was developed and applied to causal gene identification in mice and humans [11, 12]. One of the advantages of mediation analysis is that it can detect mediators at the resolution of a gene-level by leveraging the population-wide transcriptomic data. The mediator genes will then be able to help to dissect the molecular mechanisms in controlling the phenotypic variation. Indeed, by applying univariate mediation analysis in *Arabidopsis thaliana*, a study showed that SNP variants on FLOWERING TIME LOCUS C (FLC) contributed to flowering time phenotypic variation partially through different levels of FLC gene expression [13]. In plants, particularly the crop species, flowering time and other agronomically important traits are complex, possibly mediated by many mediating genes in a high dimensional manner. Therefore, the univariate mediation analysis is inadequate to tackle the complex mediation patterns.

In the present study, we leveraged the publicly available genotypic, phenotypic, and transcriptomic datasets [14]. Further, metabolomics dataset was newly generated to be used as metabolic phenotypes. We applied the recently developed high dimensional mediation analysis method [12] to these datasets and identified 736 unique and significant mediating genes controls for agronomic (N = 40) and metabolic (N = 66) traits. Noticeably, one of the identified genes, Zm00001d007718 (*Bx13*) — an essential gene in the benzoxazinoids synthesis pathway, mediates 37 phenotypes. Our results suggested that high dimensional genome-wide meditation analysis is a promising causal gene inference method to connect genotype with phenotype by integrating multi-Omics datasets.

## RESULTS

### Modeling metabolic and agronomic traits using the high-dimensional genome-wide mediation analysis

We conducted the high dimensional mediation analysis, one of the causal inference methods, to model association between genotype (exposure variable, Z) and phenotype (outcome variable, Y) by including a third hypothetical variable, gene expression, as the mediator (M) [12]. In the analysis, the assumed causal relationships are that genomic variant contributes to phenotypic variantion either directly (Z → Y) or indirectly through gene expression mediation (Z → M → Y), or both (Z → Y and Z → M → Y) (**Figure 1**).

**Fig. 1.**
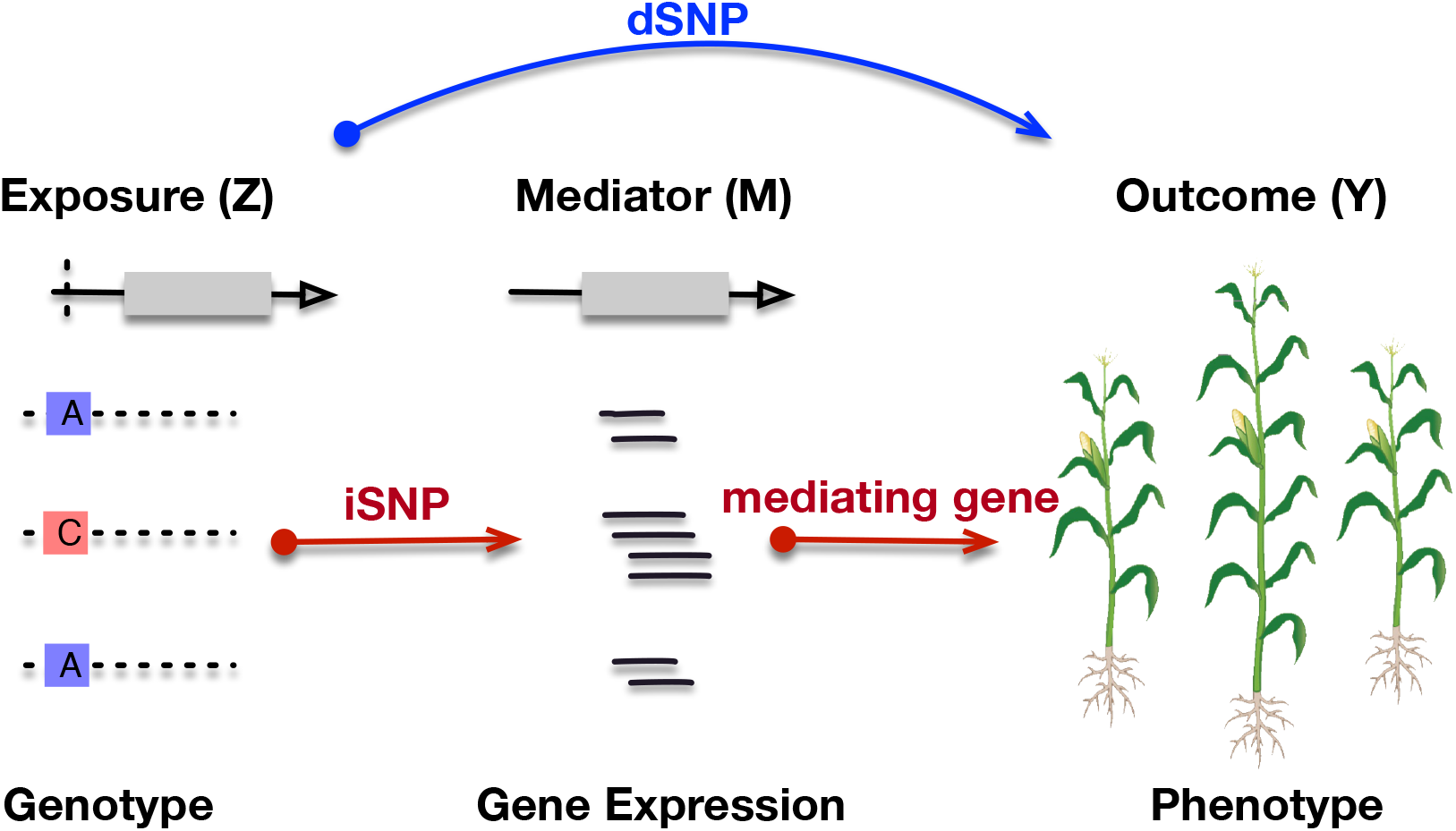
Diagram of the high-dimensional genome-wide mediation analysis. In the analysis, the SNP variant was modeled as the exposure (Z), gene expression (i.e., measured using RNA-seq read count) as a mediator (M), and plant phenotype (i.e., plant height) as an outcome (Y). The blue line with an arrow denotes the direct relationship between Z and Y through direct SNP (dSNP). The red lines with arrows denote the indirect relationship between Z and Y mediated through mediating gene that was controlled by the indirect SNP (iSNP).

To carry out the analysis, we employed the maize diveristy panel [14], because genotypic [15], phenotypic [16], and transcriptomic [17] datasets for this population were publicly available. Additionally, we collected the above-ground tissues from the two-week-old seedlings and conducted gas chromatography-mass spectrometry (GC-MS) analysis to obtain profiles of 66 primary metabolites (see **materials and methods**). Among these metabolites, the contents of 14 amino acids and their derivative N-containing metabolites showed strong positive correlations among species (mean Pearson correlation coefficient (*r*) = 0.55), while the rest of them exhibited relatively weak correlations (mean |*r*| = 0.14) (**Figure S1**). The maize diversity panel is composed of inbred lines from stiff stalk, non-stiff stalk, tropical/subtropical, sweet corn, popcorn, and mixed inbred lines (**Figure S2**) [14]. To account for the population structure, we fitted the first three principal components computed from the genome-wide markers to the mediation model as confounders (see **materials and methods**). For the analysis, we tested each normalized agronomic and metabolic trait against each of the seven RNA-seq data obtained from different tissue types as the mediators to conduct a total of N=742 (106 traits × seven tissues) analysis.

### Genome-wide mediation analysis identified direct SNPs, indirect SNPs, and mediating genes

After conducting the analysis, a set of 18,369 (9,989 unique ones) direct SNPs (dSNPs) (**Supplementary Table S1**), 24,383 (19,775 unique ones) indirect SNPs (iSNPs) (**Supplementary Table S2**), and 932 (736 unique ones) mediating genes were identified (**Supplementary Table S3**).

According to the mediation model (**Figure 1**), dSNPs were assumed to control phenotypic variation without considering gene expression mediation, much like the signals detected through GWAS. Indeed, by comparing dSNPs identified from different tissues, N=322 common dSNPs were detected regardless of the mediating tissue type (**Figure 2A**), which is significantly higher than expected by chance (efficient multi-set intersection test [18], p-value < 1.0*e* − 43). In constract, very few iSNP (N=3) and mediating gene (N=1) were detected across seven tissues (**Figure 2B, C**).

**Fig. 2.**
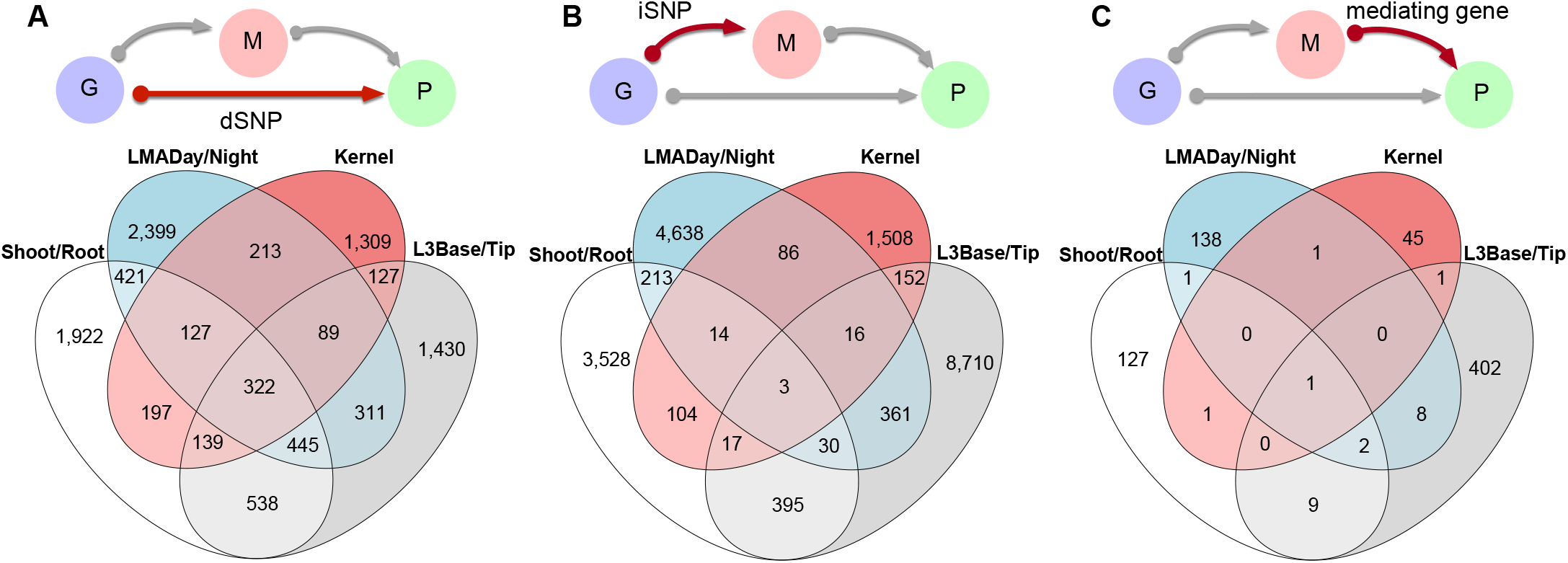
The Venn diagrams of the dSNPs, iSNPs, and mediating genes detected in different tissues. In the Venn diagrams, three pairs of tissues were grouped together based on their high relatedness. Shoot/Root, germinating shoot and root; L3Base/Tip, leaf base/tip at 3rd leaf stage; LMADay/Night, mature leaf collected during the day/night; Kernel, kernels collected 350 growing degree days after pollination.

Since the dSNPs are largely independent from the mediating tissue type, we aggregated the dSNPs identified from different tissues and find that on average 47.95% and 12.52% of the dSNPs were overlapped with the GWAS SNPs for the metabolic and agronomic traits using the window size of 5kb (**Figure S3**). At individual phenotypic trait levels, 86% and 96% of the overlaps between dSNPs and GWAS SNPs for the metabolic and agronomic trait categories were significantly higher than expected by chance (Pearson’s Chi-squared test, adjusted p-value < 0.05). The similar but slightly elevated ratios were observed with increased window sizes (**Figure S3**).

### Tissue specificity of the phenotypic variance explained by mediators

To evaluate the phenotypic variance mediated through gene expression, we calculated the PVM (*V_IE_*/(*V_IE_* + *V_DE_*)) value for each trait and RNA-seq tissue type (see **materials and methods**). For metabolite traits, gene expression data collected from the leaf tip at the 3rd leaf stage, a tissue type close to which the metabolite data were collected, exhibited significantly larger PVM (Tukey’s test, adjusted p-values < 0.01) than values calculated from other tissue types (**Figure 3**), suggesting the tissue specificity of the gene expression-based mediation model. The agronomic traits, likely regulated by more complex intermediate processes, showed significantly smaller PVM values as compared to metabolite traits in 2/7 tissue types (student t-test, p-value < 0.01), i.e., leaf 3 tip and adult leaf at night (**Figure 3**).

**Fig. 3.**
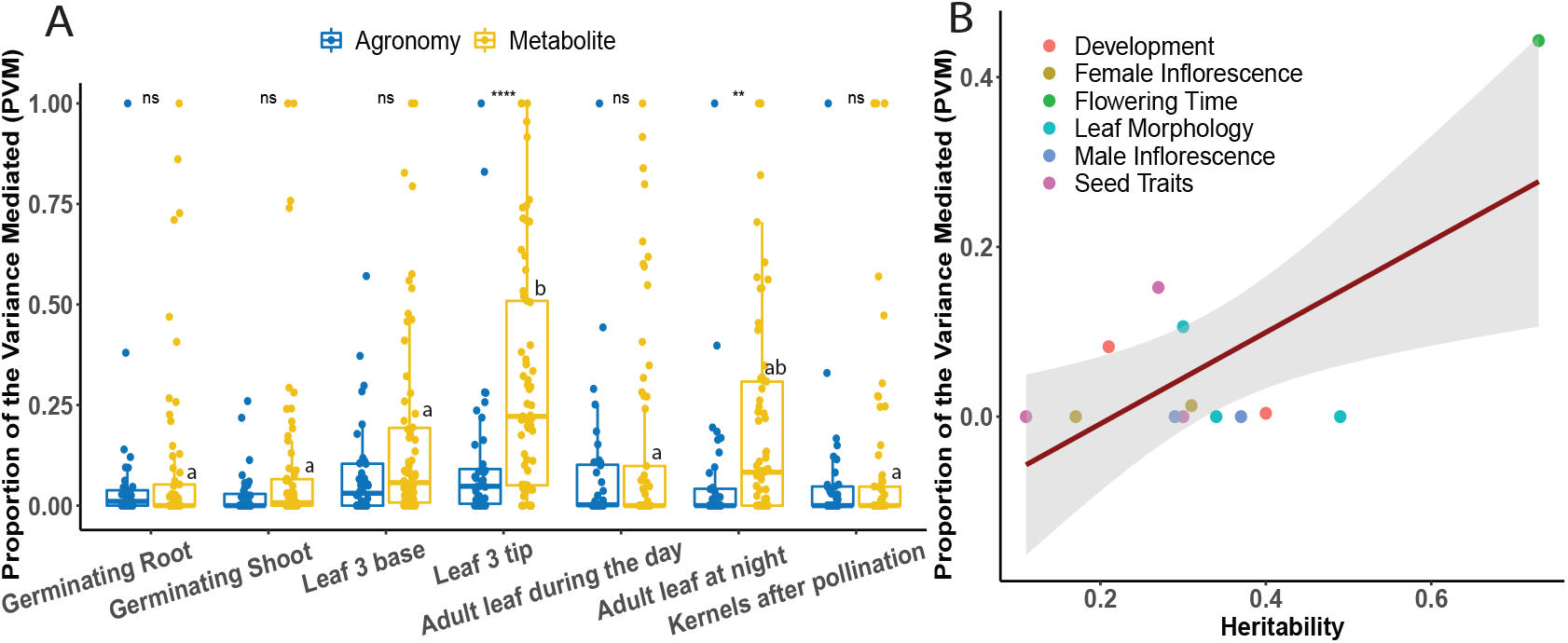
The proportion of variance mediated (PVM) by tissue type and trait category. (**A**) The PVM values for the agronomic and metabolic traits broken down by tissue type. The Tukey’s tests across seven tissues for both both agronomic and metabolic traits; ns, difference is not significant; **, p-value <0.01; and ***, p-value <0.0001. The tissue with label “a” is significantly different from tissues with label “b”, while the tissue with “ab” is not significantly different from either tissue with label a or b. (**B**) The correlation(estimate: 0.666, p-value = 0.01) between the PVM values and heritability of the agronomic traits in the adult leaf tissue collected during the day. The dashed red line denotes the regression line. The grey shaded area denotes the 95% confidence interval.

To further explore the relationship between trait heritability and the variance explained by mediators, we subdivided the agronomic traits into eight categories, including flowering time (N=6), development (N=2), female inflorescence (N=4), male inflorescence (N=2), leaf morphology (N=3), and seed traits (N=6) according to Hung et al., [16] and obtained heritability values for N=13 agronomic traits [19]. As a result, heritability was significantly correlated (*r* = 0.67, p-value = 0.013) with the PVM in the tissue of the adult leaf collected during the day (**Figure 3B**). To mitigate the possible bias driven by a flowering time trait, we conducted Fisher transformation [20] and obtained an even higher correlation coefficient (*r* = 0.80). The pattern of positive correlation between heritability and PVM, however, was not observed by modeling RNA-seq data collected from other tissue types (**Figure S4**).

### The pleiotropic mediating genes controlling multiple agronomic and metabolic traits

The N = 736 mediating genes, including 264 detected for agronomic traits and 486 for metabolic traits, were widely distributed across the maize genome. Gene ontology (GO) term enrichment test [21] found mediating genes controlling the flowering time traits were significantly enriched in response to stimulus (p-value = 2.2*e* − 3, FDR = 0.039). And mediator genes of metabolites were significantly enriched in signal transduction (p-value = 1.8*e* − 4, FDR = 0.029) and cellular development process (p-value = 1.1*e* − 4, FDR = 0.029).

We considered genes mediating for more than one trait as pleiotropic mediating genes. In total, 83/736 (11%) pleiotropic mediating genes were identified, nine of which controlled more than five traits (**Figure 4A**). Interestingly, 4/9 of these highly pleiotropic mediators mapped to the gene models with previously annotated gene functions, i.e., Zm00001d007716 — sensitive to freezing 2 (*SFR2*) [22], Zm00001d007718 — 2-oxoglutarate-dependent dioxygenase (*Bx13*) [23], Zm00001d018573 — Rho-related protein from plants 4 (*Rop4*) [24], and Zm00001d010661 — VQ motif-transcription factor43 (*VQ43*) [25, 26] (see **Supplementary Table S4** to find detailed traits mediated by these genes).

**Fig. 4.**
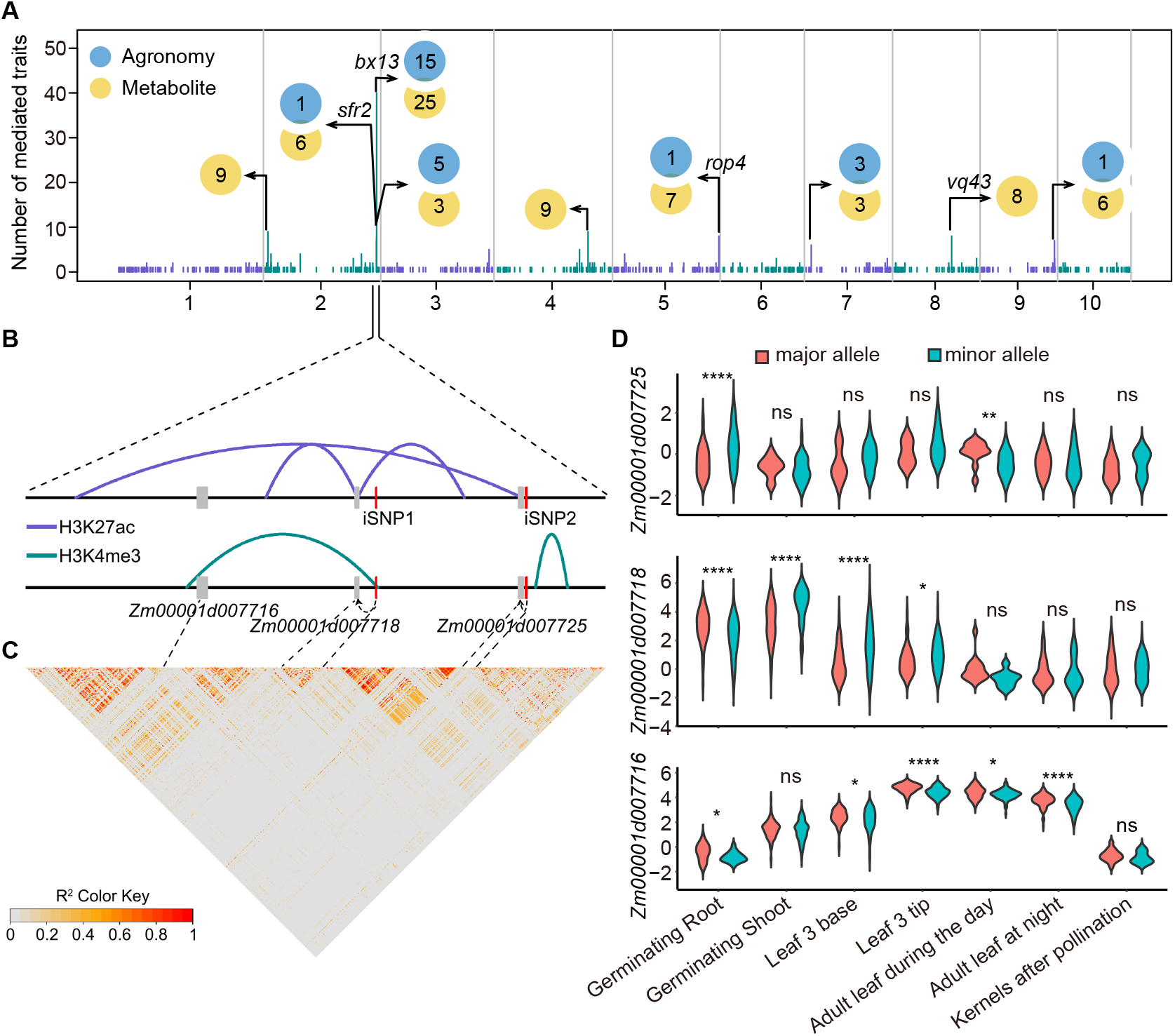
Distriution and genomic characteristics of the pleiotropic mediating genes. (**A**) Physical distribution of the pleiotropic mediating genes and the number of traits mediated by them. (**B**) Physical interactions in the chromosome 2 region. Grey boxes indicate the gene models and red lines indicate the indirect SNPs. Dashed arrowed lines denote the mediating genes controlled by the indirect SNPs. (**C**) LD block around the region. (**D**) The expression levels (unit: log(FPM), (FPM: fragments per million mapped fragments)) of the three adjacent mediating genes located on chromosome 2.

Notably, three of these highly pleiotropic genes *SFR2*, *Bx13*, and Zm00001d007725 were colocalized in a physically proximate region on chromosome two (**Figure 4B**); and they mediated 7 (1/6), 40 (15/25), and 8 (5/3) phenotypic traits (agronomy/metabolite), respectively. The low linkage disequilibrium (LD) among these three genes contradicted our initial hypothesis that these genes might be blocked in the same haplotypes (**Figure 4C**). The further study identified interactive loops around the regions using HiChIP data [27], suggesting these three pleiotropic genes might share the same regulatory modulation (**Figure 4B**). The correlation coexpression patterns of these three genes were strong in leaf base and tip at 3rd leaf stage tissues (**Fig S5**). Consistently, we detected significantly different gene expression patterns for *SFR2* and *Bx13* in leaf base and tip between lines carrying major and minor alleles (see **Figure 4D** for gene expression differences in other tissues). Taken together, these lines of evidence suggested the possibly co-regulated genes with distinct functions might mediate a number of different phenotypes.

### A pleiotropic mediator gene *Bx13* controlled 40 agronomic and metabolic traits

One of these pleiotropic mediating genes, *Bx13*, mediated 37 traits (15 agronomic and 25 metabolic traits) in tissues of the germinating shoot and leaf tips/bases at 3rd leave stage; and it was controlled by N = 598 iSNPs (**Figure 4**). The *Bx13* gene has been well characterized in a number of recent studies [23, 28, 29]. It encodes the 2-oxoglutarate-dependent dioxygenase, a key enzyme involved in benzoxazinoids metabolism [28]. According to Handrick et al., [28], the *Bx13* locus is segregating in the maize population, that an early flowering sweet corn P39 with AA genotype has higher expression than the classical temperate B73 with GG genotype in youngest leaves. Consistent with this observation, we found the maize lines carrying the minor allele with “AA” genotype at SNP *S*2_238153041 exhibited significantly (p-values = 1.7*e* − 04 and 1.9*e* − 02 in leaf base and tip at 3rd leaf stage, respectively) higher gene expression levels as opposed to the minor allele with “GG” genotype (**Figure 4D**).

Studies revealed that benzoxazinoids are a collection of naturally occurring compounds to defend against pests and disease, and they may play important roles in regulating flowering time and auxin metabolism in maize [23]. The discovery of the mediating role of *Bx13* for GDD days to silk in leaf 3 tip (**Figure 5A**) and GDD anthesis to silking interval in leaf 3 base (**Figure 5B**) validated its possible phenotypic consequences in controlling flowering time. Interestingly, a number of yield-related traits, such as total seed weight and total kernel volume, were also mediated by *Bx13* genes, suggesting that the pesticide gene may eventually affect crop yield. Additionally, the *Bx13* gene mediated 25 metabolic traits in three tissues, with the most potent mediating effect detected for the quinic acid — a compound likely accumulated for plant defense in tobacco [29] (**Figure 5C**). Using leaf 3 tip as the mediating tissue, six dSNPs, seven mediating genes, and 122 iSNPs were identified for the quinic acid trait, suggesting the complex genetic control of this compound. Additional work needs to be carried out to further dissect the relationship between *Bx13* and other *Bx13* mediated metabolites (**Figure 5D**).

**Fig. 5.**
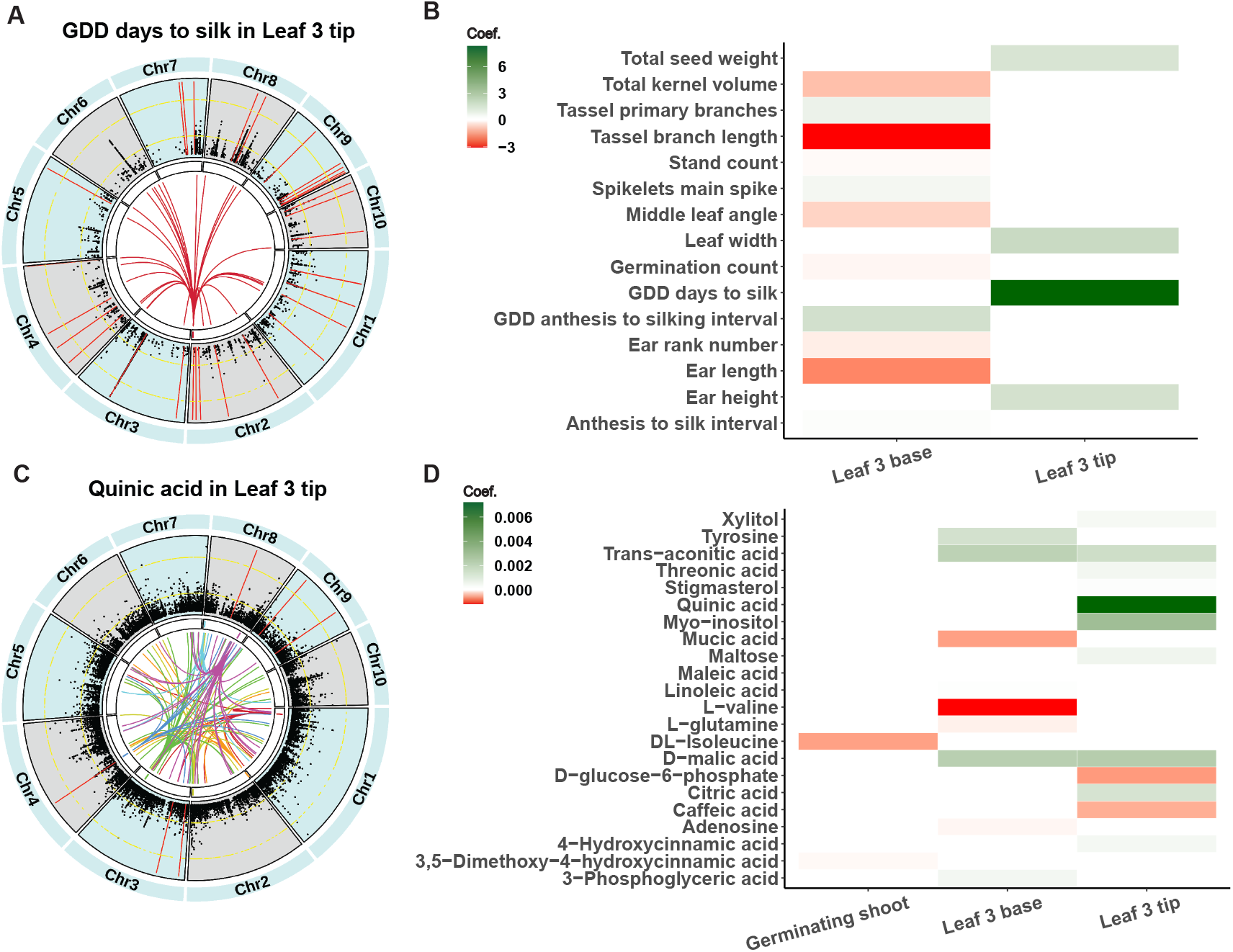
The Circos plots and the mediating effects by *Bx13* for agronomic and metabolic traits. Visualization of genome-wide mediation analysis for an agronomic trait — Growing Degree Days (GDD) to silk (**A**) and a metabolic trait — quinic acid (**C**) in the RNA-seq tissue of the leaf 3 tip. In the circos plots, the outer-most circular track represents the ten maize chromosomes; the next inner track shows the GWAS results, with two circular yellow dashed lines indicating -log(p-value) of 5 and 10 and the red lines denoting the position of direct SNPs; the next inner track shows the relative positions of identified mediating genes with different genes represented by different colors; the lines in the innermost circle connects mediators with their corresponding indirect SNPs. The heatmaps of mediating coefficients between *Bx13* gene expression and the agronomic (**B**) and metabolic traits (**D**).

## DISCUSSION

In this study, we conducted high dimensional mediation analysis and identified a number of mediating genes that were previously reported to be associated with relevant traits. Some of the identified mediating genes exhibited pleiotropic mediating effects, i.e., the *Bx13* gene mediated 15 agronomic and 25 metabolic traits. *Bx13* encodes 2-oxoglutarate-dependent dioxygenase involved in the modification of benzoxazinoids, which are defense compounds abundant in maize seedlings. The 2,4-dihydroxy-7-methoxy-1,4-benzoxazin-3-one (DIMBOA)-glucose (Gluc) and 2-hydroxy-4,7-dimethoxy-1,4-benzoxazin-3-one (HDIMBOA)-Gluc are two major benzoxazinoids in maize seedlings. BX13 enzyme mediates the first reaction to modify DIMBOA to produce other constituents of benzoxazinoids, including 2,4-dihydroxy-7,8-dimethoxy-1,4-benzoxazin-3- one (DM2BOA)-Gluc and 2-dihydroxy-4,7,8-trimethoxy-1,4-benzoxazin-3-one (HDM2BOA)-Gluc [23]. The BX13 has been repeatedly identified to be associated with the accumulation of downstream products of the pathway, i.e., DM2BOA-Gluc and HDM2BOA-Gluc [28, 30], as well as the major benzoxazinoids, DIMBOA and HDIMBOA [30, 31]. As benzoxazinoids highly accumulate in maize seedlings accounting for 1% or more of the dry weight [23], alteration in their biosynthesis likely has significant impacts on carbon and nitrogen availability in maize seedlings. Therefore, it is reasonable that the *Bx13* gene is a mediator of the accumulation of abundant metabolites in central carbon metabolism, including glutamine, trans-aconitate, and malic acid (see **Supplementary Table S4**). The results highlight the crucial effects of specialized metabolism on the central carbon metabolism. The mechanisms underlying the interactions between *Bx13* and the agronomic phenotypes need further investigation to be clarified. Benzoxazinoids are well-known to function in plant defense against various pests and pathogens [32]. Recent studies have revealed further potential functions of these compounds. For example, benzoxazinoids are shown to alter the communities of root-associated fungi and bacteria, resulting in changes in plant phenotypes [33]. Benzoxazinoids are also proposed to be involved in defense signal transduction, flowering time, growth rate, auxin signaling, and metal ion uptake [23]. Many of these postulated functions can influence phenotypes mediated by *Bx13*, including days to silking, anthesis silk interval, tassel branch length, ear length, and tassel branch length (see **Supplementary Table S4**), although precise mechanisms are still unclear.

In addition to *Bx13*, two other adjacent genes, Zm00001d007716 and Zm00001d00772, were also found mediating multiple different phenotypes. It led us to hypothesize that the three genes might be falling into the same LD block. However, LD analysis contradicted this hypothesis and suggested the three genes were segregating, although their expression levels were highly correlated in a number of tissues. The discovery of the physical contacts indicated that the three genes likely fall under similar regulation modulation in a 3-dimensional manner.

In addition to the pleiotropic mediating genes, we also identified about 500 mediating genes controlling for a single trait. The functions of some single-trait mediating genes could be verified by some relevant studies. For example, FHA-transcription factor 15 (*FHA15*, Zm00001d037971) was detected in mediating ear height (p-value = 1.1*e* − 3) and plant height (p-value = 0.23) using germinating root tissue in our study. The *FHA15* gene was associated with maize plant height and ear height in a QTL study [34]. And its possible functional role was also supported by the mutant analysis in *Arabidopsis*, where *fha15* mutant showed shortened root [35]. Moreover, Rho-related protein from plants 4 (*Rop4*, Zm00001d018573) was identified in mediating the pathogen-resistant trait in our results, consistent with the result reported previously [24].

Besides mediating genes, the high dimensional mediation analysis also detected numerous dSNPs and iSNPs. Through variance component analysis, we found the tissues with higher PVM are more likely to be responsible for traits of interest. The variance component analysis also revealed that the PVM positively correlated with the heritability of the trait, i.e., more minor variance was detected to be mediated for complex traits, at least by modeling the RNA-seq data as the mediators. For most of the traits, we only detected a limited number of mediating genes, likely due to the small sample size (N ≈ 200), the complex genetic architecture of the traits, and multi-layered intermediate molecular processes. Nevertheless, the identified mediating genes and the corresponding iSNPs paved the path for future studies to dissect the causal relationship between the genotype and phenotype via gene expression, which is critical to prioritize candidate genes to further elucidate the molecular functions for crop improvement.

## MATERIALS AND METHODS

### Public RNA-seq and phenotypic datasets for the maize diversity panel

The maize diversity panel, representing the global diversity of public maize inbreds, was chosen for the genome-wide mediation analysis [12, 14], as rich genetic and genomic resources have been accumulated over the years from the maize community. In the study, we obtained the best linear unbiased prediction (BLUP) values for N = 40 agronomic traits calculated from a multi-location multi-year field experiment [16]. The population-level transcriptomic data were obtained from Kremling et al., with RNA-seq data generated for seven different tissue types, including germinating seedling shoots/roots, leaf tips/bases at the third leaf stage, mature leaves collected during the day/night, and developing kernels [17]. The raw data contained genes with extremely low expression levels. To reduce the computational burden for the following analysis, we filtered out the genes with low expression (i.e., FPM < 1) and low variance (i.e., var < 0.05). Additionally, we filtered out genes that had a low correlation with the phenotypic trait of interest (correlation coefficient < 0.05) for the mediation analysis.

### Metabolite profiling of seedlings of the maize diversity panel

We surface-sterilized seeds of maize inbred lines from the maize diversity panel in 2% bleach for 40 minutes and imbibed them overnight in water. Seedlings were grown on pots with Fafard germination soil in a greenhouse with temperature 18-25 °C and 14/10 h light/dark cycle. We harvested the entire shoots of 14-day old seedlings within 30 min period at mid-day and quickly frozen them in liquid nitrogen. We then grounded the whole samples into fine powder under liquid nitrogen temperature and aliquoted 25 mg for metabolite analysis. We conducted metabolite profiling by using a 7200 GC-QTOF system (Agilent Technologies, Santa Clara, CA, USA) following the procedure described previously [36]. Briefly, we extracted the metabolites using methanol:water:chloroform and dried the 50 µl of the upper aqueous phase down to be derivatized by methoxyamination and trimethylsilylation. We conducted peak annotation and quantification using MassHunter Unknown and Quantitation software, respectively, with extensive manual curation. The peak retention time and the m/z of the quantitation ion are available in **Supplementary Table S5**. Finally, we calculated the relative levels of metabolites by normalizing the peak height ion count by that of internal standard (ribitol) and the sample’s precise fresh weight.

### Genotypic data processing and LD pruning

We downloaded the maize HapMap V3.2.1 (with imputation, AGPv4) genotypic data at Panzea database (https://www.panzea.org/genotypes) [15]. Using the PLINK software [37], we merged the variants on different chromosomes and retained the bi-allelic SNPs only. We then performed filtration of SNPs by discarding variants with missing rate > 0.3 across lines and a minor allele frequency (MAF) cutoff of < 0.05, which gave us a subset of 22.5 million SNPs. To reduce the SNP density and computational time in mediation analysis, we performed LD pruning of SNPs by calculating LD between each pair of SNPs in the window of 10 kb and removed one of a pair of SNPs if the LD was greater than 0.1. We then shifted the window 10 bp forward and repeated the procedure until the end of genome and resulted in a final subset of 0.77 million SNPs.

### High dimensional mediation analysis

For a population composed of *n* inbred lines, assume each of them contains *p* genes with expression data and has been genotyped for *q* SNPs. We fitted each gene separately with gene expression level as the response variable in the mediator model, the first several PCs as covariates, and all the SNPs as explanatory variables. The model is shown as below:

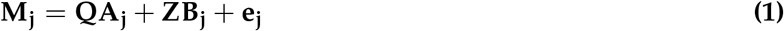

where, **M_j_**(a n×1 vector) represents the expression level of the *j*th gene; **Q** (a *n* × *s* matrix) is the first *s* PCs (*s* = 3 in our analysis); **A_j_**(a *s* × 1 vector) is the coefficients of the first *s* PCs to the *j*th gene; **Z** (a *n* × *q* matrix) represents the whole SNP set of the population composed of *n* individuals with *q* number of SNPs; *B_j_* (a *q* × 1 vector) is the coefficients of the *q*th SNPs to the *j*th gene (*j* = 1,…, *p*); and *e_j_* is the vector the residual errors with 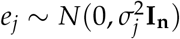.

Additionally, we fitted all the SNPs and the mediating genes in the outcome model, which uses phenotype as the response variable, as shown below.

In the model,

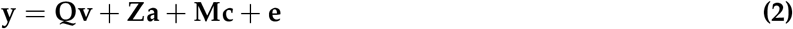

where, **y**(a *n* × 1 vector) represents the phenotype (here we used the BLUP value of a given trait); **Q** (a *n* × *s* matrix) and **Z** (a *n* × *q* matrix) are the same matrices as model **(1)**; **v**(a *s* × 1 vector) is the coefficients of first *s* PCs to phenotype; **a**(a *q* × 1 vector) denotes the coefficients of the SNPs to phenotype; **M** (a *n* × *p* matrix) is the expression levels of the *p* genes (i.e., a matrix combines all the *M_j_*); **c**(a *p* × 1 vector) is the coefficients of gene expressions to phenotype; **e** is the vector the residual errors with **e** ∼ *N*(0, *σ*^2^ *I_n_*).

### Proportion of the variance mediated

To evaluate the relative contribution of different factors to phenotypic variance, we quantified the variance due to different components as described in Qi’s study [12]. Here, we partitioned the variance into two major components, i.e., variance due to direct effect (*V_DE_*, the variance caused by effect on the change of phenotype completely due to DNA variation but not through gene expression mediation) and variance due to indirect effect (*V_ID_*, the variance caused by effect on the change of phenotype mediated by gene expression which in turn caused by DNA variation).

Using the similar notations as above formulas, we calculated the variance components using the equations as below.

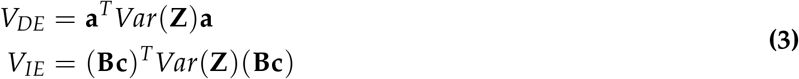

And then, we obtained the proportion of the variance mediated (PVM) as below.

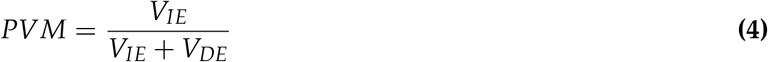

### Genome wide association study

We conducted GWAS using the GEMMA software [38] with 22.5 million SNPs. In the analysis, we fitted the first three principle components (PCs) as covariates calculated from genome-wide SNPs using the PLINK software. To account for the genetic relatedness, we fitted the centered identical by state kinship matrix generated by TASSEL software [39] as the random effects in the GWAS model.

### GO term enrichment analysis

We carried out gene ontology (GO) term analysis using agriGO v2.0 [21]. In the analysis, we employed Fisher’s exact test to perform the statistical test and Yekutieli (FDR under dependency) method to adjust multi-test problem. We used the following parameters in the analysis: minimum number of mapping = 3, gene ontology type = plant GO slim, background = locus ID v3.30 (Gramene Release 50).

## CODE AVAILABILITY

The code used for the analyses can be accessed through GitHub (https://github.com/ZhikaiYang/mediation).

## ACKNOWLEDGEMENTS

J.Y. is supported by the Agriculture and Food Research Initiative Grant number 2019-67013-29167 from the USDA National Institute of Food and Agriculture and the University of Nebraska-Lincoln Start-up fund. J.Y. and T.O. is supported by the Agriculture and Food Research Initiative Grant number 2021-67013-33898 from the USDA National Institute of Food and Agriculture and the National Science Foundation under award number OIA-1557417 for Center for Root and Rhizobiome Innovation (CRRI). Q.Z. was supported by the National Science Foundation under award numbers DBI-1564621 and OIA-1736192 when he was faculty at University of Nebraska Lincoln, and is currently supported by University of New Hampshire Start-up. This work was conducted using the Holland Computing Center of the University of Nebraska-Lincoln Start-up, which receives supports from the Nebraska Research Initiative.

## AUTHOR CONTRIBUTIONS STATEMENT

J.Y. designed this work. J.Y. and T.O. generated the data. Z. Y., G.X., Q. Z., T.O., and J.Y. analyzed the data. Z. Y. and J.Y. wrote the manuscript.

## SUPPORTING INFORMATION SUPPLEMENTAL TABLES

**Table S1.** Direct SNPs identified through genome-wide mediation analysis. (https://github.com/ZhikaiYang/mediation/blob/master/data/supplementary/Supplemental_Table_S1.csv)

**Table S2.** Indirect SNPs identified through genome-wide mediation analysis. (https://github.com/ZhikaiYang/mediation/blob/master/data/supplementary/Supplemental_Table_S2.csv)

**Table S3.** The mediating genes identified through genome-wide mediation analysis. (https://github.com/ZhikaiYang/mediation/blob/master/data/supplementary/Supplemental_Table_S3.csv)

**Table S4.** The traits mediated by pleiotropic mediating genes identified through genome-wide mediation analysis. (https://github.com/ZhikaiYang/mediation/blob/master/data/supplementary/Supplemental_Table_S4.csv)

**Table S5.** Information of the peaks annotated to the metabolites. The metabolite name, retention time, retention index, and the ion m/z used for quantification are shown for each peak annotated to a metabolite. Metabolites were quantified using the split modes indicated on the “Split Mode” column. (https://github.com/ZhikaiYang/mediation/blob/master/data/supplementary/Supplemental_Table_S5.csv)

## SUPPLEMENTAL FIGURES

**Fig. S1.**
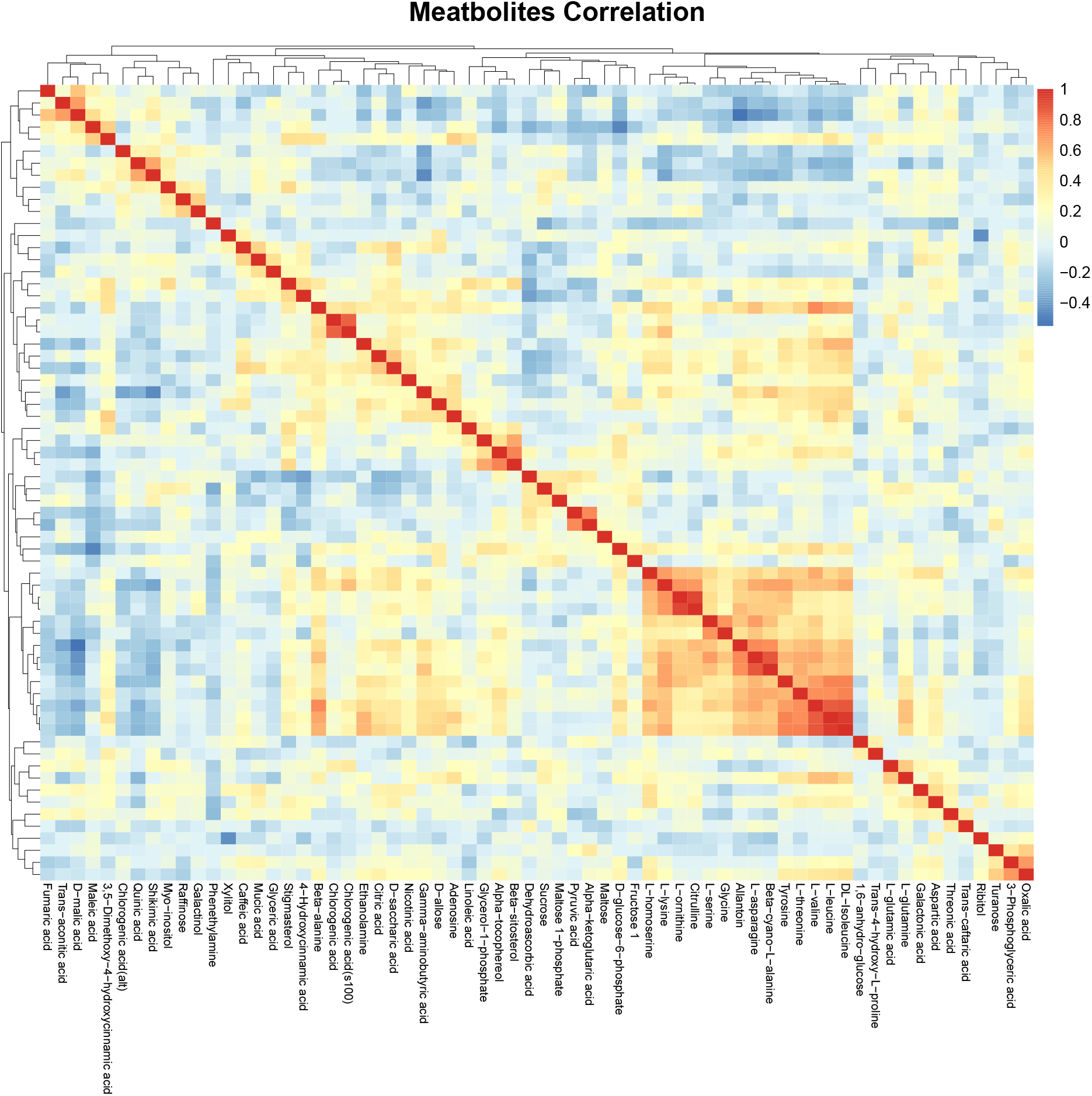
Heatmap of the correlation among N=66 metabolites. Colors denote the Pearson correlation coefficients.

**Fig. S2.**
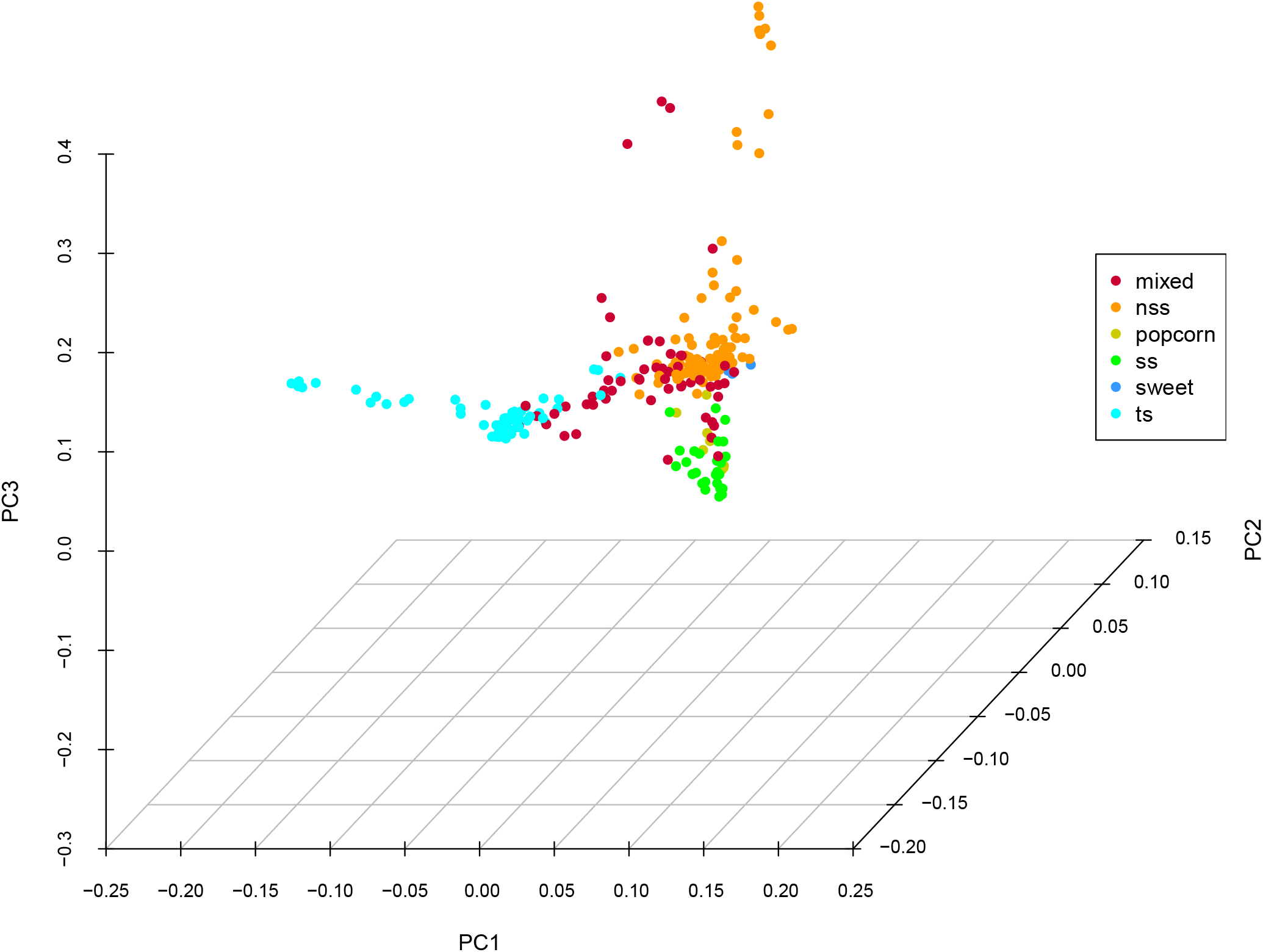
The principle component analysis of the maize population. In the legend, “mixed” denotes the mixed inbreds; “nss” denotes Non-Stiff Stalk; “popcorn” denotes pop-corn; “ss” denotes Stiff Stalk; and “sweet” denotes sweet corn; “ts” denotes Tropical/Subtropical lines.

**Fig. S3.**
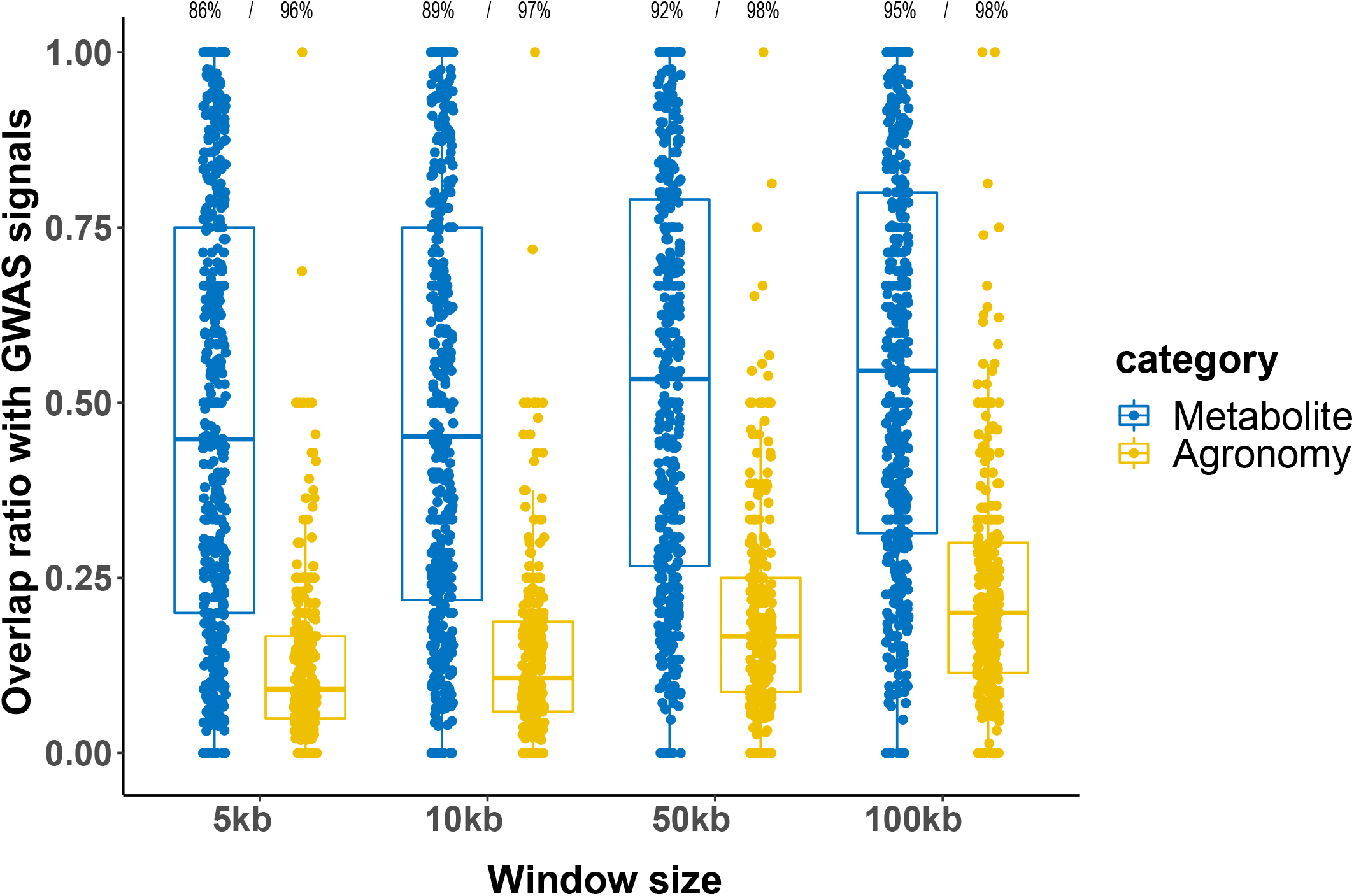
Comparison between direct SNPs and GWAS signals. The overlap ratio between direct SNP and GWAS signals for agronomic and metabolic traits. The value on top of each bar chart denotes the percentage of significant tests out of all Chi-squared tests (p-value < 0.05).

**Fig. S4.**
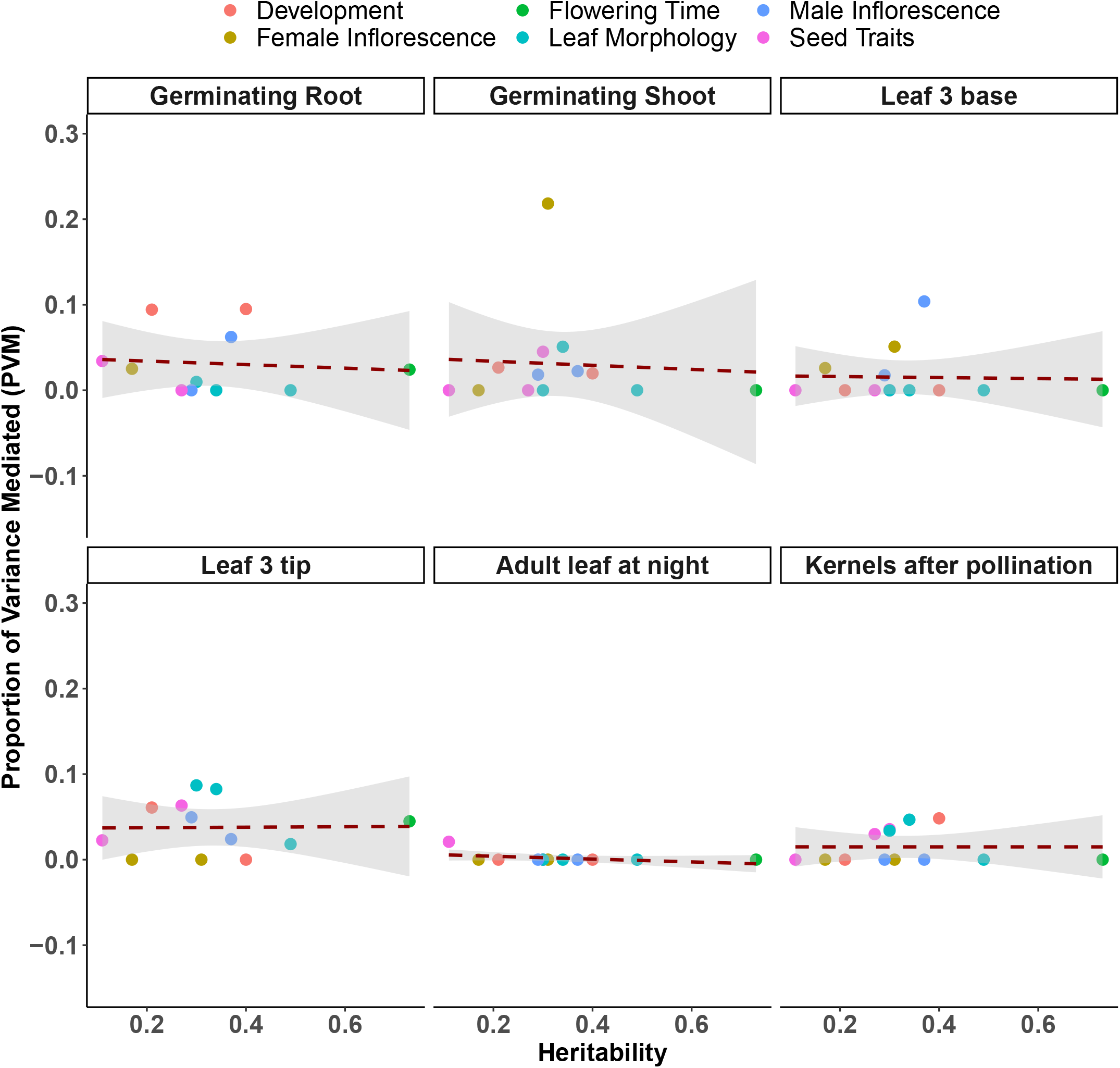
The correlation between PVM values and heritability of the agronomic traits in six tissues. The dashed lines denote the fitted line using a linear model with grey shaded areas denote the 95% confidence intervals.

**Fig. S5.**
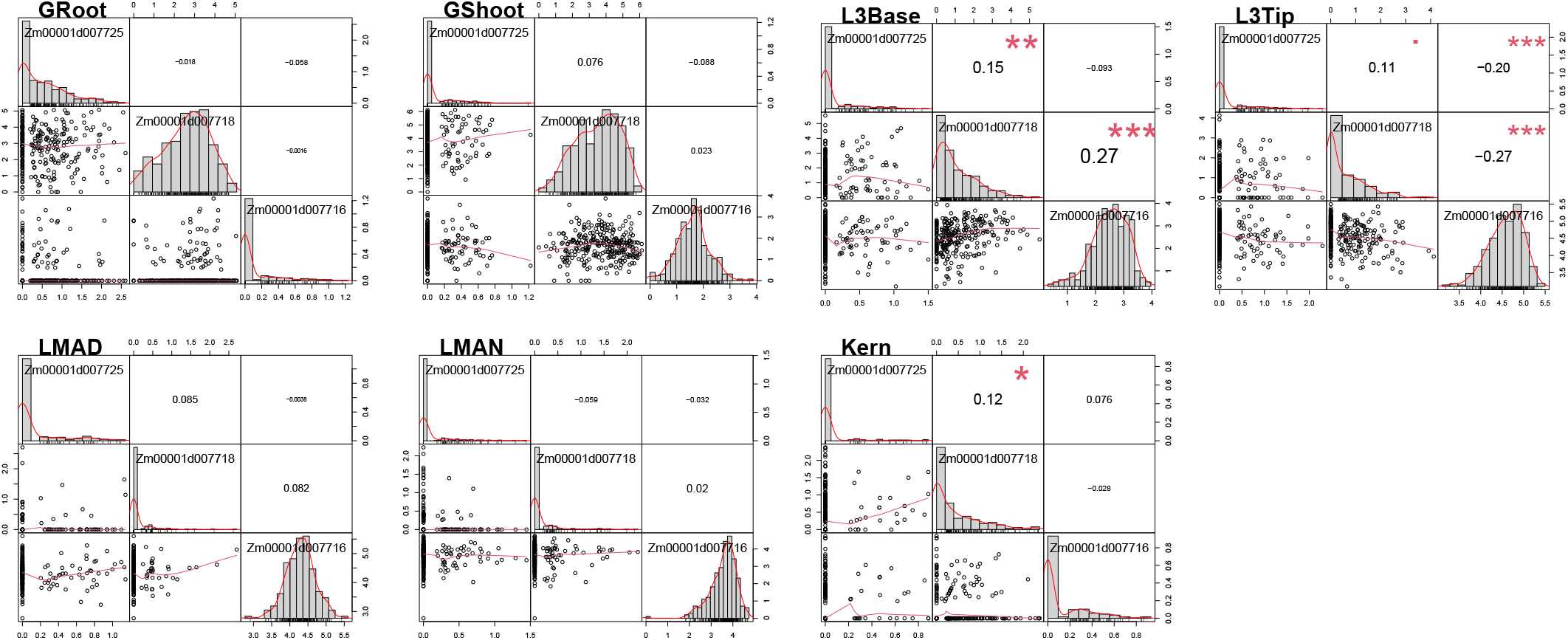
The pairwise correlations for gene expression levels of Zm00001d007716 (sfr2), Zm00001d007718 (bx13), and Zm00001d00772 in seven tissues. For each plot, the upper diagonal shows the Pearson correlation coefficient and lower diagonal shows the scatter plots of the two genes’ expression levels. On the diagonal shows the histogram of the gene expression distribution. Asterisk denotes the statistical significance with *, p-value < 0.05, **, p-value < 0.01, and ***, p-value < 0.01.

